# Comparative transcriptomics reveal differential gene expression in *Plasmodium vivax* geographical isolates and implications on erythrocyte invasion mechanisms

**DOI:** 10.1101/2023.02.16.528793

**Authors:** Daniel Kepple, Colby T. Ford, Jonathan Williams, Beka Abagero, Shaoyu Li, Jean Popovici, Delenasaw Yewhalaw, Eugenia Lo

## Abstract

*Plasmodium vivax* uses Duffy binding protein (PvDBP1) to bind to the Duffy Antigen-Chemokine Receptor (DARC) to invade human erythrocytes. Individuals who lack DARC expression (Duffy-negative) are thought to be resistance to *P. vivax*. In recent years, *P. vivax* malaria is becoming more prevalent in Africa with a portion of these cases detected in Duffy-negatives. Apart from DBP1, members of the reticulocyte binding protein (RBP) and tryptophan-rich antigen (TRAg) families may also play a role in erythrocyte invasion. While the transcriptomes of the Southeast Asian and South American *P. vivax* are well documented, the gene expression profile of *P. vivax* in Africa and more specifically the expression level of several erythrocyte binding gene candidates as compared to DBP1 are largely unknown. This paper characterized the first *P. vivax* transcriptome in Africa and compared with those from the Southeast Asian and South American isolates. The expression of 4,404 gene transcripts belong to 12 functional groups including 43 specific erythrocyte binding gene candidates were examined. Overall, there were 10-26% differences in the gene expression profile amongst the geographical isolates, with the Ethiopian and Cambodian *P. vivax* being most similar. Majority of the gene transcripts involved in protein transportation, housekeeping, and host interaction were highly transcribed in the Ethiopian *P. vivax*. Erythrocyte binding genes including *PvRBP2a* and *PvRBP3* expressed six-fold higher than *PvDBP1and* 60-fold higher than *PvEBP/DBP2*. Other genes including *PvRBP1a, PvMSP3.8, PvMSP3.9, PvTRAG2, PvTRAG14*, and *PvTRAG22* also showed relatively high expression. Differential expression was observed among geographical isolates, e.g., *PvDBP1* and *PvEBP/DBP2* were highly expressed in the Cambodian but not the Brazilian and Ethiopian isolates, whereas *PvRBP*2a and *PvRBP*2b showed higher expression in the Ethiopian and Cambodian than the Brazilian isolates. Compared to *Pvs*25, the standard biomarker for detecting female gametocytes, *PvAP2-G* (PVP01_1440800), GAP (PVP01_1403000), and *Pvs47* (PVP01_1208000) were highly expressed across geographical samples. These findings provide an important baseline for future comparisons of *P. vivax* transcriptomes from Duffy-negative infections and highlight potential biomarkers for improved gametocyte detection.

## 1. Introduction

*Plasmodium vivax* Duffy binding protein (PvDBP1), which binds to the cysteine-rich region II of the human glycoprotein Duffy Antigen-Chemokine Receptor (DARC) (1–3), was previously thought to be the exclusive invasion mechanism for *P. vivax* (4). However, reports of *P. vivax* infections in Duffy-negative individuals (3) have raised important questions of how *P. vivax* invades erythrocytes that lack DARC expression. It was hypothesized that either mutations in *PvDBP1* or a weakened expression of DARC allowed *P. vivax* to invade Duffy-negative erythrocytes (5, 6). Despite several mutational differences observed in *PvDBP1* between Duffy-positive and Duffy-negative infections, these differences do not lead to binding of Duffy-negative erythrocytes (4) and suggested an alternative invasion pathway.

The *P. vivax* nuclear genome is ~29 megabases with 6,642 genes distributed amongst 14 chromosomes (7). Remarkably, across the *P. vivax* genome, approximately 77% of genes are orthologous to *P. falciparum, P. knowlesi*, and *P. yoelii* (8). Genes involved in key metabolic pathways, housekeeping functions, and membrane transporters are highly conserved between *P. vivax* and *P. falciparum* (8). However, at the genome level, *P. vivax* isolates from Africa, Southeast Asia, South America, and Pacific Oceania are significantly more polymorphic than the *P. falciparum* ones (9, 10), likely due to differences in distributional range, transmission intensity, frequency of gene flow via human movement, and host susceptibility (11).

In *P. vivax*, erythrocyte binding protein (*PvEBP*), reticulocyte binding protein (*PvRBP*), merozoite surface protein (*PvMSP*), apical membrane antigen 1 *(PvAMA1)*, anchored micronemal antigen (*PvGAMA*), Rhoptry neck protein (*PvRON*), and tryptophan-rich antigen genes (*PvTRAg*) families have been suggested to play a role in erythrocyte invasion (9, 12).

*PvDBP1, PvMSP1, PvMSP7*, and *PvRBP2c* were previously shown to be highly polymorphic (13–17). *PvEBP*, a paralog of *PvDBP1*, harbors the hallmarks of a *Plasmodium* red blood cell invasion protein and is similar to *PcyM DBP*2 sequences in *P. cynomolgi* that contains a Duffy-binding like domain (18). Binding assay of *PvEBP* region II (171-484) showed moderate binding to Duffy-negative erythrocytes (4). Both *PvDBP*1 and *PvEBP* (*PvEBP/DBP*2 hereafter) exhibit high genetic diversity and are common antibody binding targets associated with clinical protection (19, 20). Several members of *PvRBP2 (PvRBP2a, PvRBP2b, PvRBP2c, PvRBP2d, PvRBP2e, PvRBP2p1*, and *PvRBP2p2)* are orthologous to *PfRh2a, PcyRBP2*, and *PfRh5*, with *PvRBP*2a and *PfRh*5 share high structural similarity (21, 22). *PvRBP*2b and *PvRBP*2c are orthologous to *PcyRBP2b* and *PcyRBP2c*, respectively (23). The receptor for PvRBP2a was previously identified as CD98, a type II transmembrane protein that links to one of several L-type amino acid transporters (24); the receptor for PvRBP2b is transferrin receptor 1 (TfR1) (25). The PvRBP2b-TfR1 interaction plays a critical role in reticulocyte invasion in Duffy-positive infections (25). MSP1 also shows a strong binding affinity, with high-activity binding peptides (HABPs) clustered close to fragments at positions 280–719 and 1060–1599 (26), suggesting a critical role in erythrocyte invasion. Although the *MSP*7 gene family shows no binding potential, it forms a complex with *PvTRAg36.6* and *PvTRAg56.2* on the surface, likely for stabilization at the merozoite surface (27). A comparison of *P. vivax* transcriptomes between *Aotus* and *Saimiri* monkeys indicated that the expression of six *PvTRAg* genes in *Saimiri P. vivax* was 37-fold higher than in the *Aotus* monkey strains (28), five of which bind to human erythrocytes (27, 29). Although most PvTRAg receptors remain poorly characterized, the receptor of PvTRAg38 has been identified as Band 3 (30).

Recent progress in transcriptomic sequencing of *P. vivax* in non-human primates has provided an overview of stage-specific gene expression profile and structure, of which thousands of splices and unannotated untranslated regions were characterized (31, 32) The transcriptomes of Cambodian (33) and Brazilian (34) *P. vivax* field isolates showed high expression levels and large populational variation amongst host-interaction transcripts. Heterogeneity of gene expression has been documented amongst *P falciparum-* infected samples, implying that the parasites can modulate the gene transcription process through epigenetic regulation (35). However, the transcriptomic profile of African *P. vivax* remains unexplored, and it is unclear if there is heterogeneity among the geographical isolates. In addition, our previous study found that two CPW-WPC genes PVP01_0904300 and PVP01_1119500 expressed in the male gametocytes, and *Pvs230* (PVP01_0415800) and *ULG8* (PVP01_1452800) expressed in the female gametocytes were highly expressed relative to *Pvs25* in the Ethiopian *P. vivax* (36). While these genes have a potential to be used for gametocyte detection, it remains unclear if such expressional patterns are similar in other geographical isolates.

In this study, we aimed to 1) examine the overall gene expression profile of 10 Ethiopian *P. vivax* with respect to different intraerythrocytic lifecycle stages; 2) determine the expression levels of previously characterized erythrocyte binding gene candidates (9); 3) compare gene expression profiles of the Ethiopian *P. vivax* with the Cambodian (33) and Brazilian (34) isolates from *in vitro* especially for the erythrocyte binding and male/female gametocyte gene candidates. These findings provide the first description of the *P. vivax* transcriptomes in Africa. A systematic comparison of gene expression profiles among the African, Southeast Asian, and South American isolates will deepen our understanding of *P. vivax* transcriptional machinery and invasion mechanisms.

## 2. Materials and Methods

### 2.1 Ethics statement and data availability

Scientific and ethical clearance was obtained from the institutional scientific and ethical review boards of Jimma University, Ethiopia and University of North Carolina, Charlotte, USA. Written informed consent/assent for study participation was obtained from all consenting heads of households, parents/guardians (for minors under 18 years old), and individuals who were willing to participate in the study. Sequences for the 10 Ethiopian transcriptomes are available on the National Center for Biotechnology Information Short Read Archive under BioProject: PRJNA784582. All code is available on GitHub at https://github.com/colbyford/vivax_transcriptome_comparisons.

### 2.2 Sample preparation

Ten microscopy-confirmed *P. vivax* samples were collected from Duffy positive patients at hospitals in Jimma, Ethiopia. These patients had 4,000 parasites/μL parasitemia and had not received prior antimalarial treatment. A total of 10mL whole blood was preserved in sodium heparin tubes at the time of collection. Red blood cell pellets were isolated and cryo-preserved with two times glycerolyte 57 and stored in liquid nitrogen. Prior to culture, samples were thawed by adding 0.2V of 12% NaCl solution drop-by-drop followed by a 5-minute room temperature incubation. Ten-times volume of 1.6% NaCl solution was then added drop-by-drop to the mixture and the samples were centrifuged at 1000 rcf for 10 minutes to isolate the red blood cell pellet. This process was repeated with a 10x volume of 0.9% NaCl. Following centrifugation, the supernatant was removed via aspiration, and 18mL of sterile IMDM (also containing 2.5% human AB plasma, 2.5% HEPES buffer, 2% hypoxanthine, 0.25% albumax, and 0.2% gentamycin) per 1mL cryo mixture was added to each sample for a final hematocrit of 2%. 10% Giemsa thick microscopy slides were made to determine the majority parasite stage and duration of incubation required, averaging 20-22 hours for the majority trophozoites and 40-44 hours for the majority ring. Samples were incubated at 37°C in a 5% O2, 5% CO2 atmosphere to allow growth to the schizont stage. *In vitro* maturation was validated through microscopic smears 20-40 hours after the initial starting time, dependent on the majority stage (Supplementary Figure 1A). To minimize oxidative stress, each culture was checked more than two times and returned to a 5% oxygen environment immediately after checking.

Cultured pellets were isolated via centrifugation and placed in 10x volume trizol for RNA extraction. RNA extraction was performed using direct-zol RNA prep kit according to the manufacturer’s protocol, followed by two rounds of DNA digestion using the DNA-free kit (Zymo). Samples were analyzed with a nanodrop 2000 and RNA Qubit to ensure sample concentrations were above 150 ng total for library construction. For samples with no significant amount of DNA or protein contaminants, RNA libraries were constructed using Illumina rRNA depletion library kits according to the manufacturer’s protocol. Completed libraries were quality checked using a bioanalyzer to ensure adequate cDNA was produced before sequencing. Sample reads were obtained using Illumina HiSeq 2×150bp configuration to obtain at least 35 million reads per sample. Sequence reads were aligned with HISAT2 (37), using the Rhisat2 R package (38) to the P01 *P. vivax* reference genome and all human reads were filtered out using SAMtools (39) (implemented in the R package (40)). The alignment was mapped to the P01 reference annotation using the Rsubread package (41).

### 2.3 Data analyses

To further confirm samples were majority schizont stage, sequence reads of each sample were deconvoluted in CIBERSORTx (42) based on *P. berghei* homologs (43). We used the published matrix to determine the frequency of expression for each gene calculated for rings, trophozoites, and schizonts, respectively. Transcripts that were expressed 30% or more were sorted into their respective stages (Supplementary Figure 1B). All reads were annotated using the Rsubread package and classified into 12 different categories by function. We then examined the top 30 transcribed genes using the counts per million (CPM) metric.

Our previously published whole genome sequence data identified several mutations and structural polymorphisms in genes from the *PvEBP, PvRBP, PvMSP*, and *PvTRAg* gene families that are likely to involve in erythrocyte invasion (9). Specific binding regions in some of the genes such as *PvDBP*1, *PvEBP/DBP2*, *PvRBP*2b, and *PvMSP*3 have been identified (44). To further explore the putative function, we compared relative expression levels of 43 erythrocyte binding gene candidates (Supplementary Table 1) in the 10 Ethiopian *P. vivax* samples with other geographical isolates that were of majority schizont stage. We used the CPM and TPM (transcripts per million) metrics in R package edgeR (45). The CPM metric was used to obtain the top 30 transcripts overall and does not consider gene length, while TPM considers gene length for normalization and allows an unbiased conclusion to be made relative between and to other transcriptomes (34). We then transformed the data using *log*(2)TPM+1 to illustrate relative expression levels via a heat map with an average abundance. We also selected 25 gametocyte gene candidates, 15 of which were shown to correlate to female gametocyte development and nine to male gametocytes (36, 46), to assess their expression levels relative to the standard *Pvs25* in the samples. In addition, we examined the expression of AP2-G that is a critical transcription factor for both male and female gametocyte development (47).

### 2.4 Comparison of datasets

RNA-seq data of four *in vitro* Cambodian (33) and two *in vitro* Brazilian (34) *P. vivax* samples were downloaded from the GitHub repository and analyzed with the same bioinformatic methods described above to minimize potential batch effects. Samples were deconvoluted using the same matrix. The Ethiopian *P. vivax* samples were cultured and sequenced using similar protocol as the Cambodian (33) and Brazilian (34) ones with slight modifications in media and library preparation. We obtained the average expression and standard deviation in TPM for each gene target and determined potential difference in transcription levels by conducting pairwise differential expression (DE) analysis among the Cambodian, Brazilian, and Ethiopian samples. The expression level of 6,829 genes were examined for DE by edgeR dream (45, 48) and variancePartition (49), with adjusted *p*-value<1.0e-6 for DE gene concordance. A linear mixed effects models was used to ensure accuracy in triplicated Brazilian samples, and the Kenward-Roger method was used to estimate the effective degree of freedom for hypothesis testing due to small sample sizes.

## 3. Results

### 3.1 Overview of the Ethiopian *P. vivax* transcriptomes

All 10 Ethiopian *P. vivax* samples originated from Duffy-positive patients. Based on deconvolution, all 10 Ethiopian *P. vivax* samples had similar proportions of trophozoite and schizont stage (Figure 1A). Only less than 1% of the sequence reads belong to the ring stage.

**Figure 1.**
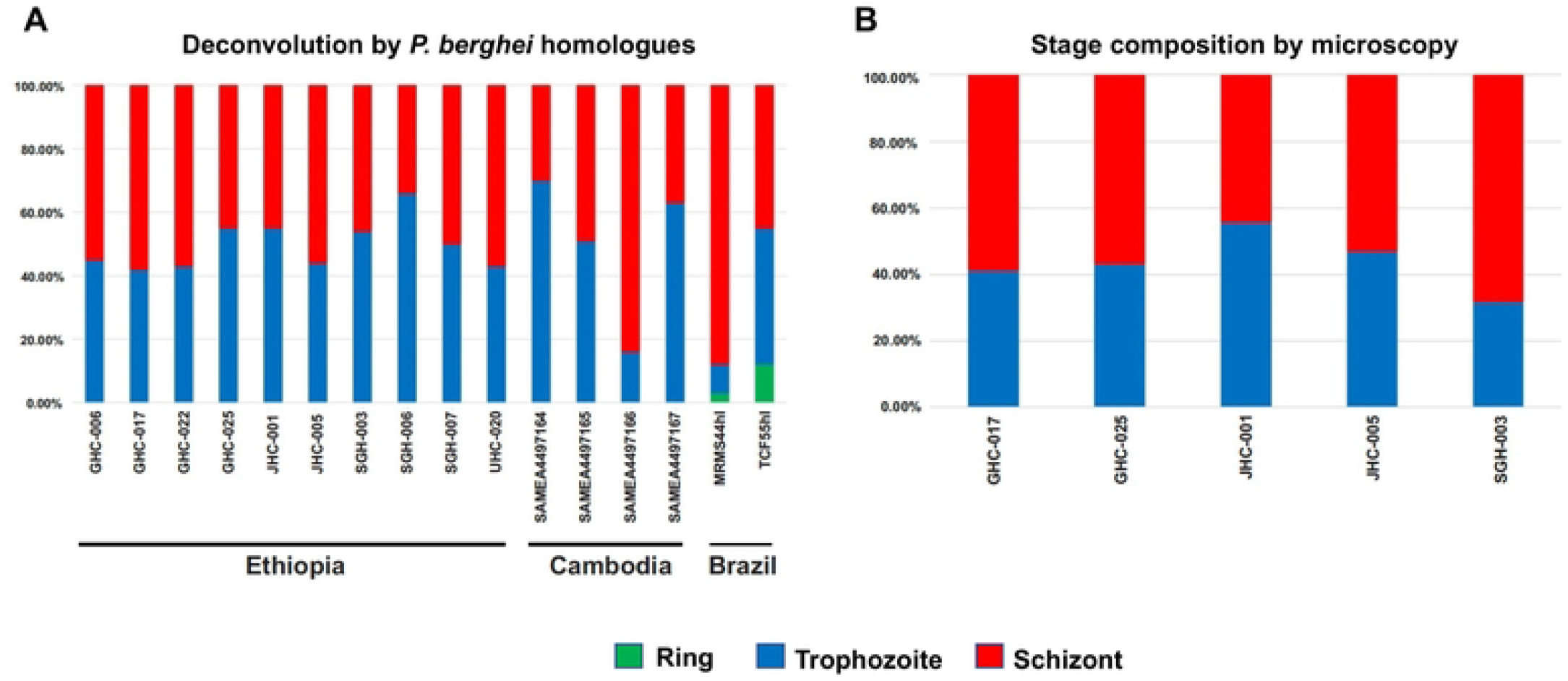
(A) CIBERSORTx deconvolution of the 10 Ethiopian, four Cambodian, and two Brazilian *P. vivax* transcriptomes using a *P. berghei* homologue matrix. No significant difference was observed in the proportion of trophozoites and schizonts amongst the isolates (*p*>0.05). (B) Parasite stage based on microscopic analysis of five Ethiopian *P. vivax* samples. No significant difference was observed between microscopy and computational deconvolution for these samples (*p*>0.05).

Microscopic results corroborated the deconvolution analyses showing similar proportion of parasite stages in a subset of samples (Figure 1B). The deconvolution of *P. vivax* sequence reads from the Cambodian and Brazilian samples also showed no significant difference in the proportions of trophozoites or schizonts (*P*>0.05; Figure 1A).

Overall, about 64% (4,404 out of 6,830) of the genes were detected with transcription in the Ethiopian *P. vivax* (Supplementary Table 2). Of the 4,404 genes, 69% (2,997) were annotated with known functions and 31% (1,407 genes) remain uncharacterized (Figure 2A). We normalized each sample expression profile to TPM to remove technical bias in the sequences and ensure gene expressions were directly comparable within and between samples (Supplementary Table 2). Of the 2,997 genes with known function, 21.7% are responsible for housekeeping, and 14.2% genes for post-translation modifications (PTMs) and regulation. The PIR proteins account for 4.8% (212) of all the identified genes and ~2.8% of the genes are involved in host-pathogen interactions. Nearly 52% of all detectable transcripts (2,288 genes) were expressed at a threshold of 20 TPM or above, which were considered as highly transcribed (Figure 2B). These highly transcribed transcripts showed similar proportions of gene categories including unknown, PTM/regulatory, DNA regulation, replication/elongation, host interactions, cell signaling, and resistance. Only transcripts involved in transport and housekeeping showed a slight increase of 2.9% and 1.48%, respectively, indicating a higher activity relative to the other categories. By contrast, transcripts involved in RNA regulation, PIR, and ribosomal activity showed a slight decrease of 2.19%, 1.79%, and 1.71%, indicating an overall lower activity compared to other categories (Figure 2B).

**Figure 2.**
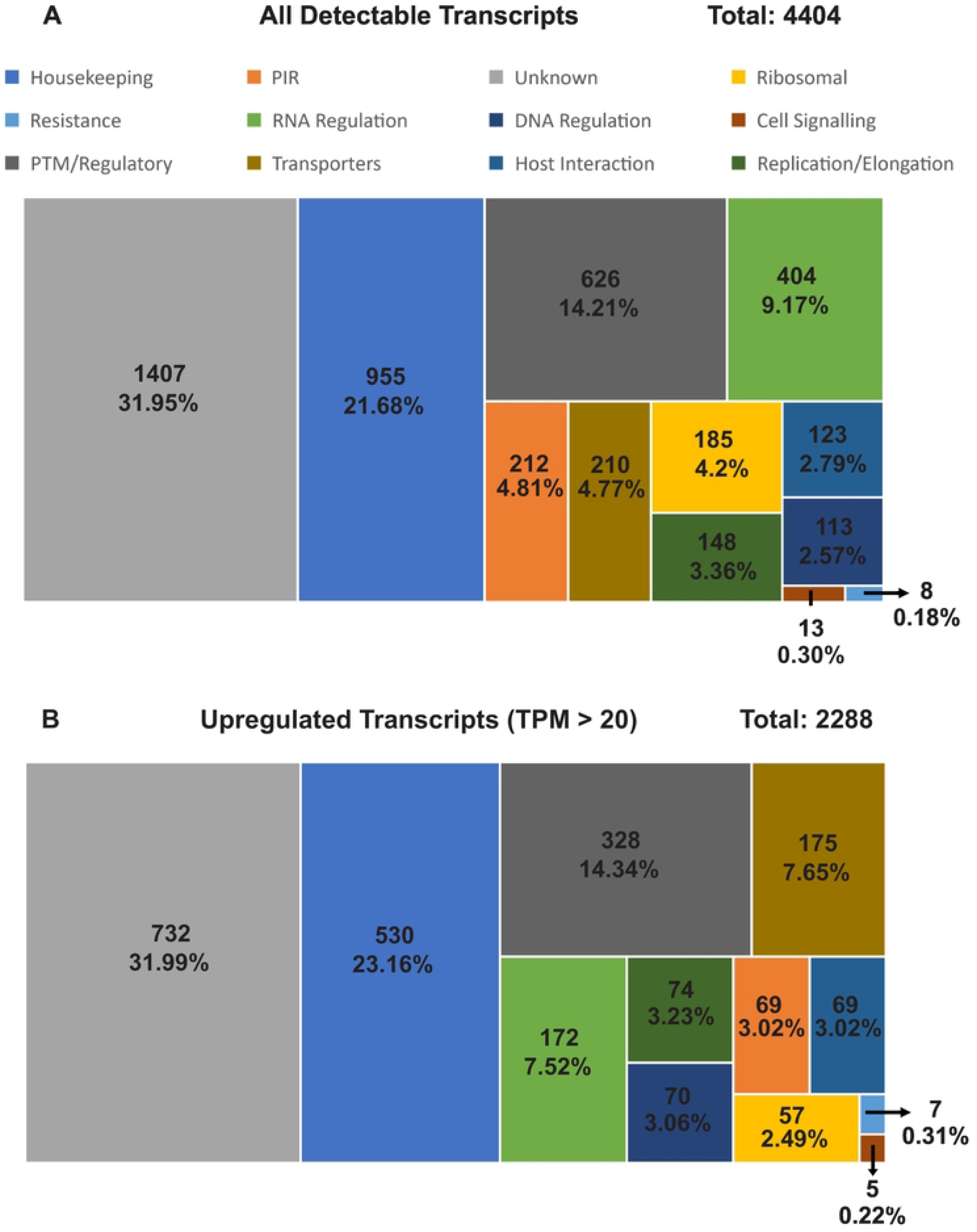
Categorization of (A) all detectable transcripts and (B) upregulated (TPM > 20) transcripts for the Ethiopian *P. vivax* by gene function. The numbers shown represent the number of transcripts along with the overall percentage compared to all detected transcripts. Transcripts that were not detected were removed from the analysis. Only transcripts involved in transport and housekeeping showed a slight increase of 2.9% and 1.48%, respectively in the number of upregulated transcripts, indicating a higher activity relative to the other categories. By contrast, transcripts involved in RNA regulation, PIR, and ribosomal activity showed a slight decrease of 2.19%, 1.79%, and 1.71%, indicating an overall lower activity compared to other categories.

### 3.2 Top 30 highly expressed transcripts of Ethiopian *P. vivax*

For the 10 Ethiopian *P. vivax* transcriptomes, four genes including PVP01_1000200 (PIR protein), PVP01_0202900 (18s rRNA), PVP01_0319600 (RNA-binding protein), and PVP01_0319500 (unknown function) were the most highly expressed among the others (Figure 3). Transcripts involved in housekeeping and PTM regulation each account for 23.3% of the top 30 highly expressed genes. Among genes involved in host-interactions, PVP01_0715400 (merozoite organizing protein), PVP01_0816800 (protein RIPR), PVP01_1402400 (reticulocyte binding protein 2a), and PVP01_1469400 (reticulocyte binding protein 3) are highly expressed. Five gene transcripts including PVP01_1000200 from the PIR family, PVP01_0319500 of unknown function, PVP01_0202900 a 18S rRNA, PVP01_1329600 a putative glutathione S-transferase, and PVP01_0418800 a putative pentafunctional AROM polypeptide showed most variable expression levels among the 10 samples, with a standard deviation of 20,000 and higher CPM (Figure 3). Three other genes including PVP01_0202700 (28S ribosomal RNA), PVP01_1137600 (basal complex transmembrane protein 1), PVP01_1243600 (replication factor C subunit 3) showed moderate variation ranging from 1,397 to 1,033 CPM. All other genes such as PVP01_1206500 (elongation factor Tu) and PVP01_1011500 (an unclassified protein) showed consistent expression level with variation under 1,000 CPM among samples (Figure 3).

**Figure 3.**
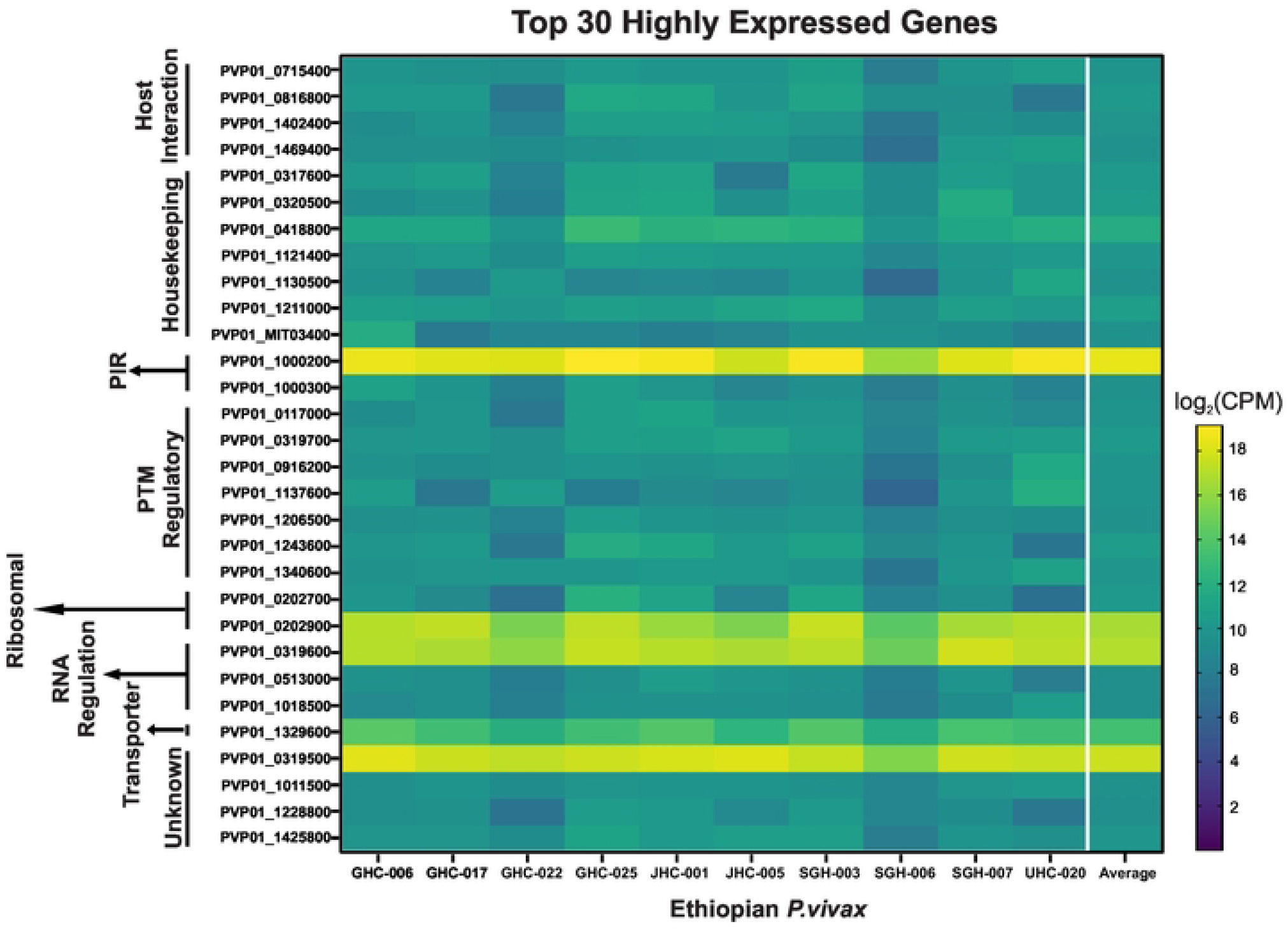
Heat map showing the top 30 highly transcribed genes based on *log*(2)CPM+1. Genes are arranged by different functions as indicated on the y-axis. Overall, four genes including PVP01_1000200 (PIR protein), PVP01_0202900 (18s rRNA), PVP01_0319600 (RNA-binding protein), and PVP01_0319500 (unknown function) from four different functional groups were shown to be most highly expressed among the others. Of interest, PVP01_0715400 (merozoite organizing protein), PVP01_0816800 (protein RIPR), PVP01_1402400 (reticulocyte binding protein 2a), and PVP01_1469400 (reticulocyte binding protein 3) were among the top 30 highly expressed genes involved in host interactions.

### 3.3 Differentially expressed genes among geographical *P. vivax*

The overall gene expression profile was similar between the Ethiopian and Cambodian *P. vivax*, but different from the Brazilian ones (Figure 4A; Supplementary Table 3). Several genes involved in DNA regulation, host-interactions, replication, ribosomal, and transportation were upregulated in the Ethiopian and Cambodian isolates but showed considerable downregulation in Brazilian ones. Based on the Kenward-Roger DE analyses, a total of 1,831 differentially expressed genes were detected between the Cambodian and Brazilian isolates (CvB), 1,716 between the Ethiopian and Brazilian (EvB), and 721 between the Ethiopian and Cambodian (EvC) isolates (Figure 4B-D). The EvC analysis showed the lowest differentiation with only 10.6% of the entire transcriptome (Figure 4B), while EvB and CvB showed a greater differentiation of 25.1% and 26.8%, respectively (Figures 4C & D). For the 721 genes that were differentially expressed between the Cambodian and Ethiopian *P. vivax*, nearly half of them were significantly upregulated in Ethiopia compared to Cambodia (Figure 4B). Four genes including PVP01_0208700 (V-type proton ATPase subunit C), PVP01_0102800 (chitinase), PVP01_0404000 (PIR protein), and PVP01_0808300 (zinc finger (CCCH type protein) showed low levels of transcription (log_10_*P*-value>50; Figure 4B) compared to other DE genes. By contrast, two genes including PVP01_1329600 (glutathione S-transferase) and PVP01_MIT03400 (cytochrome b) were highly transcribed (log_2_fold change>10). For the 1,716 genes that were differentially expressed between the Ethiopian and Brazilian *P. vivax*, 914 of them were highly transcribed (Figure 3C). Of these, three genes including PVP01_1412800 (M1-family alanyl aminopeptidase), PVP01_0723900 (protein phosphatase-beta), and PVP01_0504500 (28S ribosomal RNA) showed a log_10_*P*-value greater then 75, indicating substantial expressional differences. For the 1,831 genes that were differentially expressed between the Cambodian and Brazilian *P. vivax*, 948 of them were highly transcribed (Figure 4D). Four genes including PVP01_1005900 (ATP-dependent RNA helicase DDX41), PVP01_0318700 (tRNAHis guanylyltransferase), PVP01_1334600 (60S ribosomal protein L10), and PVP01_1125300 (SURP domain-containing protein) showed substantial expressional differences with log_10_*P*-value greater than 75. Two genes, PVP01_0010550 (28S ribosomal RNA) and PVP01_0422600 (early transcribed membrane protein), were shown with low expression (*log*_10_fold change<-12), while one gene PVP01_0901000 (PIR protein) with substantial expression (*log*_10_fold change>12). These comparisons further demonstrated the differences in transcriptional patterns between geographical isolates.

**Figure 4.**
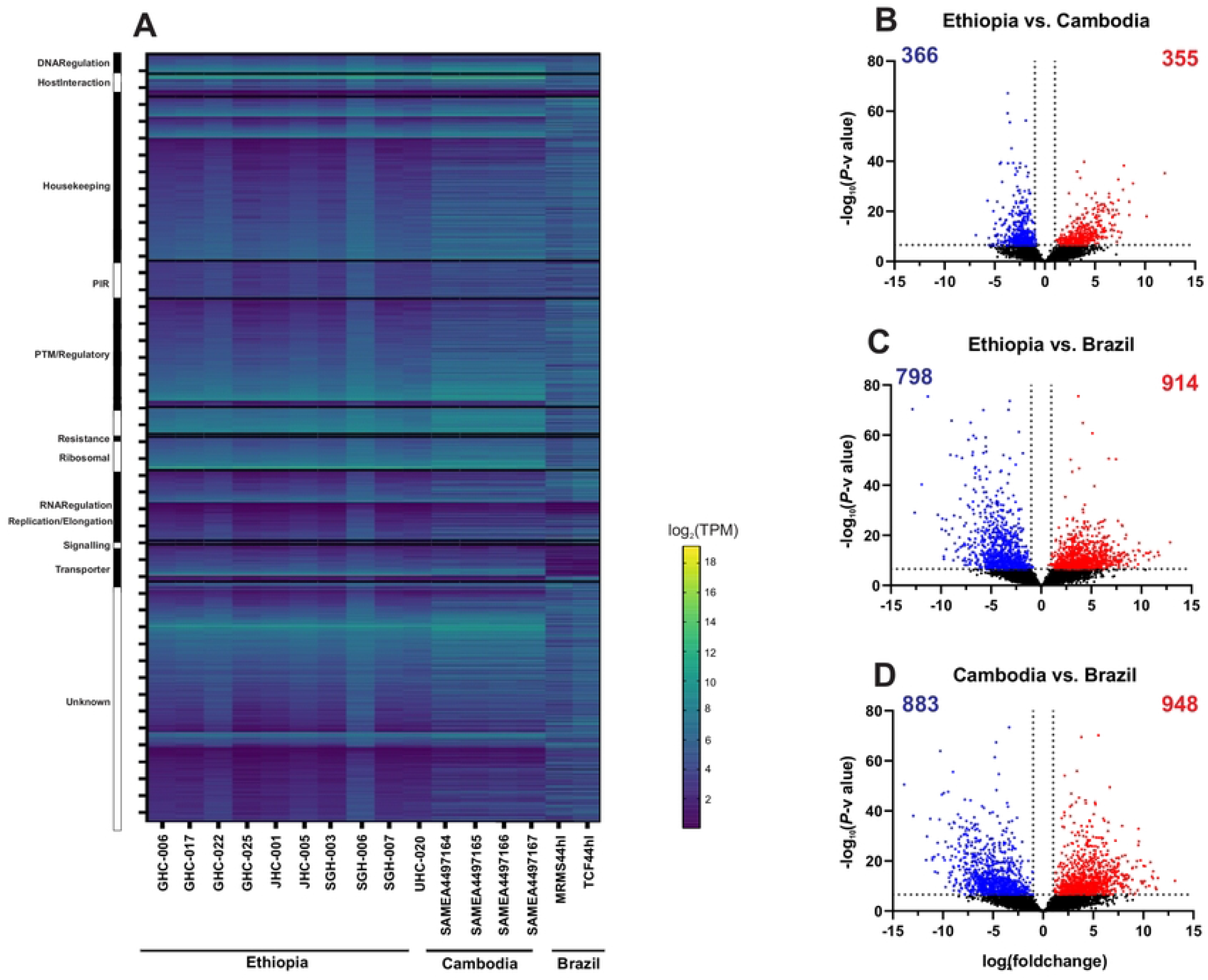
(A) Comparisons of the entire transcriptomes with genes sorted by functionality among the Ethiopian, Cambodian, and Brazilian *P. vivax*. The overall gene expression profile was nearly identical between the Ethiopian and Cambodian *P. vivax*, but different from the Brazilian isolates. Several genes involved in DNA regulation, host-interactions, replication, ribosomal, and transportation were upregulated in the Ethiopian and Cambodian isolates but showed considerable downregulation in Brazilian ones. (B-D) Volcano plots based on the Kenward-Roger DE analyses comparing differentially expressed genes between the (B) Ethiopian and Cambodian; (C) Ethiopian and Brazilian; (D) Cambodian and Brazilian isolates. Blue dots represent single genes that are downregulated in the comparison while red dots represent upregulated genes by comparison. About 10% of the detectable transcripts were differentially expressed between the Ethiopian and Cambodian *P. vivax*, but about 25% and 27% variations were detected between the Ethiopian and Brazilian as well as the Cambodian and Brazilian *P. vivax*, respectively. Overall, the Brazilian isolates had more genes that were upregulated compared to the Ethiopian and Cambodian ones.

### 3.4 Expression of genes related to erythrocyte invasion

Of the 43 candidate genes associated with erythrocyte binding function, *PvDBP1* on average showed about 10-fold higher expression than *PvEBP/DBP*2, which showed very low expression in four of the Ethiopian *P. vivax* samples (Figure 5). *PvRBP2b* showed four-fold higher expression than *PvEBP/DBP2*, but 50% less than *PvDBP1. PvRBP2a* showed consistently the highest expression across all samples, with about 6-fold, 67-fold, and 15-fold higher expression than *PvDBP1, PvEBP/DBP2*, and *PvRBP2b*, respectively. Other genes including *PvMSP*3.8, *PvTRAg14*, and *PvTRAg22* also showed higher expression than *PvDBP1*. Of the 15 *PvTRAg* genes, only *PvTRAg14* and *PvTRAg22* showed expression higher than *PvDBP1; PvTRAg23* and *PvTRAg*24 showed the lowest expression. Other putatively functional ligands including *PvRA* and *PvRON4* showed 7-10 times lower expression compared to *PvDBP1*, though *PvGAMA, PvRhopH3, PvAMA1*, and *PvRON2* were expressed higher than *PvEBP/DBP2*.

**Figure 5.**
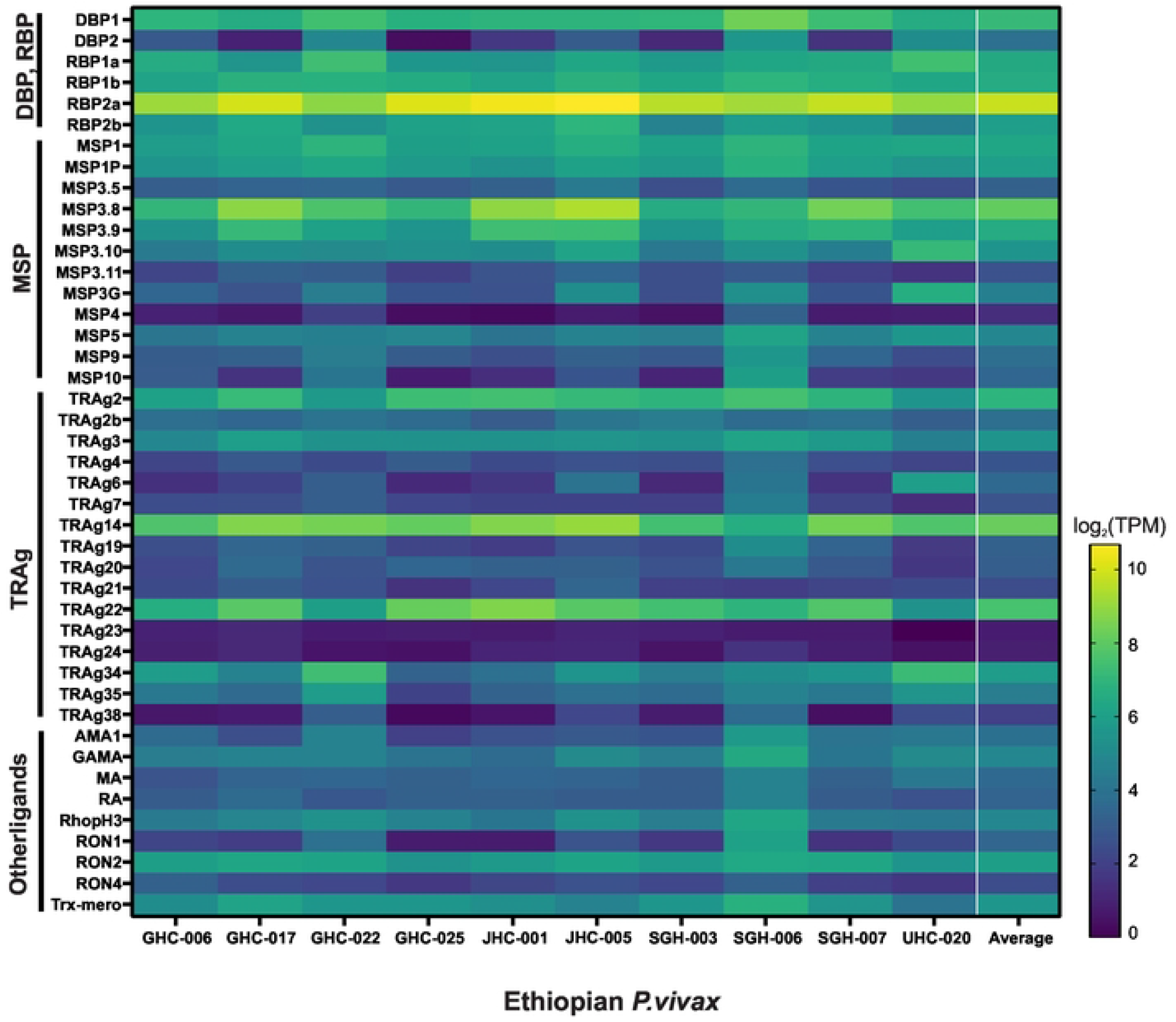
Heatmap showing 43 genes associated with erythrocyte binding function in the Ethiopian *P. vivax* based on *log*(2)TPM+1 values. *PvRBP2b* showed four-fold higher expression on average than *PvEBP/DBP2*, but 50% less than *PvDBP1. PvRBP2a* showed consistently the highest expression across all samples, with about 6-fold, 67-fold, and 15-fold higher expression than *PvDBP*1 *, PvEBP/DBP*2, and *PvRBP*2b, respectively. Other genes including *PvMSP*3.8, *PvTRAg*14, and *PvTRAg22* also showed higher expression than *PvDBP*1.

We further compared the expressional pattern of these 43 genes in the Ethiopian *P. vivax* with the Cambodian and Brazilian isolates (Figure 6). Members of the *PvDBP* and *PvRBP* gene family showed generally higher expression in the Cambodian *P. vivax* than the other isolates (Figure 6A). For instance, the expression of *PvDBP1, PvRBP1a*, and *PvRBP1b* were significantly higher in the Cambodian than the other isolates (*P*<0.01), whereas *PvRBP2a* and *PvRBP*2b showed higher expression in the Ethiopian *P. vivax* than the others. Compared to the *PvDBP* and *PvRBP* gene families, the expression patterns of *PvMSP* were different (Figure 6B).

**Figure 6.**
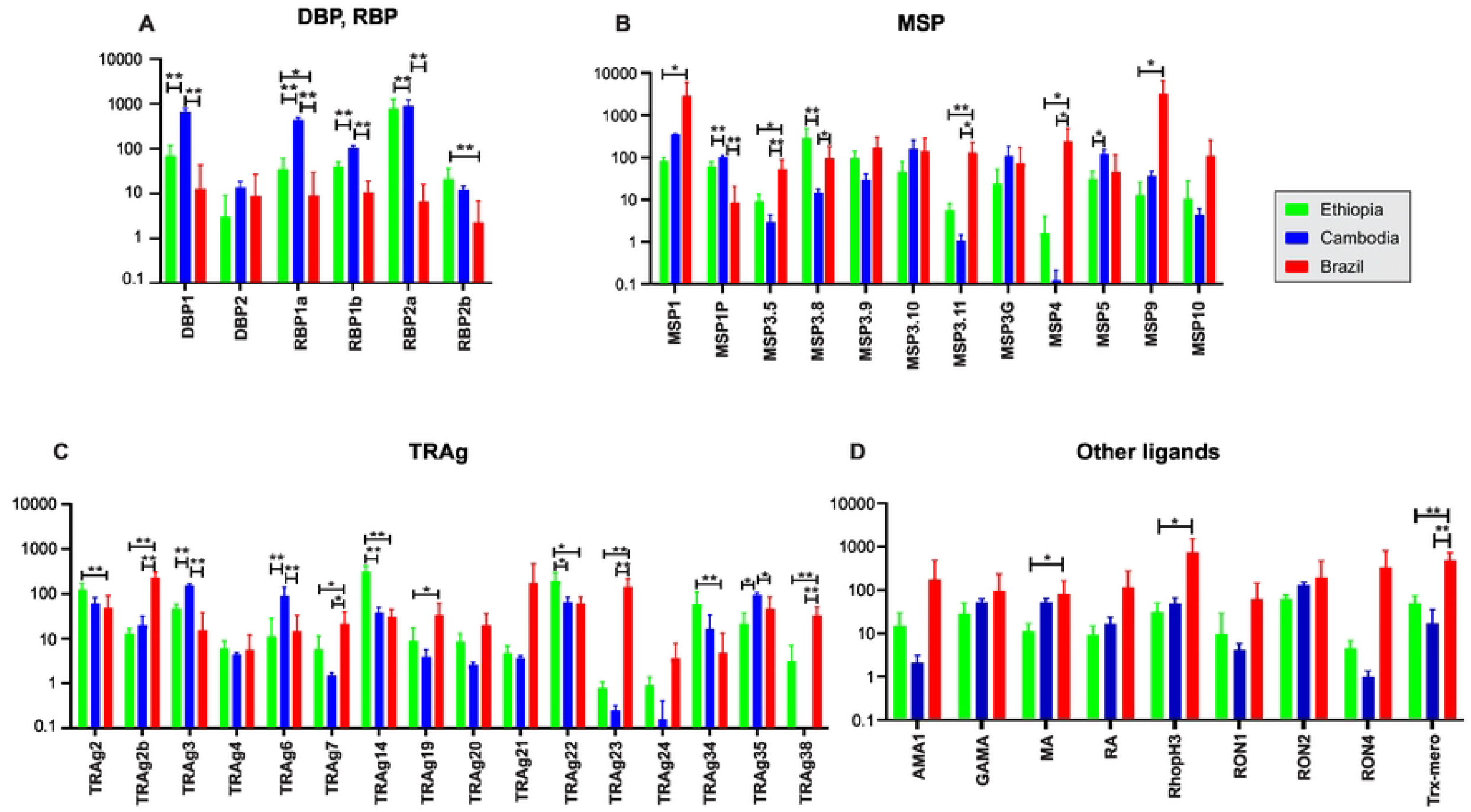
Comparisons of 43 genes associated with erythrocyte binding function based on *log*(2)TPM+1 values across the Ethiopian, Cambodian, and Brazilian *P. vivax* for (A) *PvDBP1, PvEBP*, and *PvRBP* genes; (B) *PvMSP* genes; (C) *PvTRAg* genes; (D) other putatively functional ligands. * denotes *P*-value <0.05; ** denote *P*-value <0.01.

Most of the *MSP* gene members including *PvMSP3.5, PvMSP3.11*, and *PvMSP4* showed substantially higher expression in the Brazilian *P. vivax* than the other isolates (*P*<0.01). Only *PvMSP3.8* of the 12 *PvMSP* genes was expressed significantly higher in the Ethiopian than the others (*P*<0.01; Figure 6B). Of the 16 *PvTRAg* genes, *PvTRAg14* and *PvTRAg22* showed significantly higher expression in the Ethiopian isolates compared to the others (*P*<0.05; Figure 6C). Eight other members including *PvTRAg*2b, *PvTRAg*7, *PvTRAg*19, *PvTRAg*20, *PvTRAg*21, *PvTRAg23, PvTRAg24*, and *PvTRAg38* showed significantly higher expression in the Brazilian isolates than the others (*P*<0.05; Figure 6C). The remaining nine putatively functional ligands showed relatively similar expression levels, except for *PvMA, PvRhopH3*, and *PvTrx-mero* that were highly expressed in the Brazilian isolates (*P*<0.05; Figure 6D).

### 3.5 Expression of female and male gametocyte genes

Based on the expression level of *Pvs25* (PvP01_0616100), all 10 Ethiopian *P. vivax* samples contained submicroscopic gametocytes, in addition to the four samples from Cambodia and two samples from Brazil (Figure 7). Amongst the 26 gametocyte-related genes, *PvAP2-G* (PVP01_1440800) as well as the gametocyte associated protein, GAP (PVP01_1403000) and *Pvs47* (PVP01_1208000) from female and male gametocytes, respectively, showed the highest expression across the Ethiopian, Cambodian, and Brazilian isolates, and were consistently higher than *Pvs*25 (Figure 7). This expression pattern suggests the potential utility of these three genes as better gametocyte biomarkers across geographical isolates. Other genes indicated differential expression patterns among isolates, e.g., the female gametocyte gene PVP01_0904300 (CPW-WPC family protein) showed consistently high levels of expression in both the Ethiopian and Cambodian isolates, though much lower in the Brazilian ones. On the other hand, PVP01_1302200 (high mobility group protein B1) and PVP01_1262200 (fructose 1,6-bisphosphate aldolase) from the female and male gametocytes showed the highest expression levels in Brazilian *P. vivax* but not the Ethiopian and Cambodian ones.

**Figure 7.**
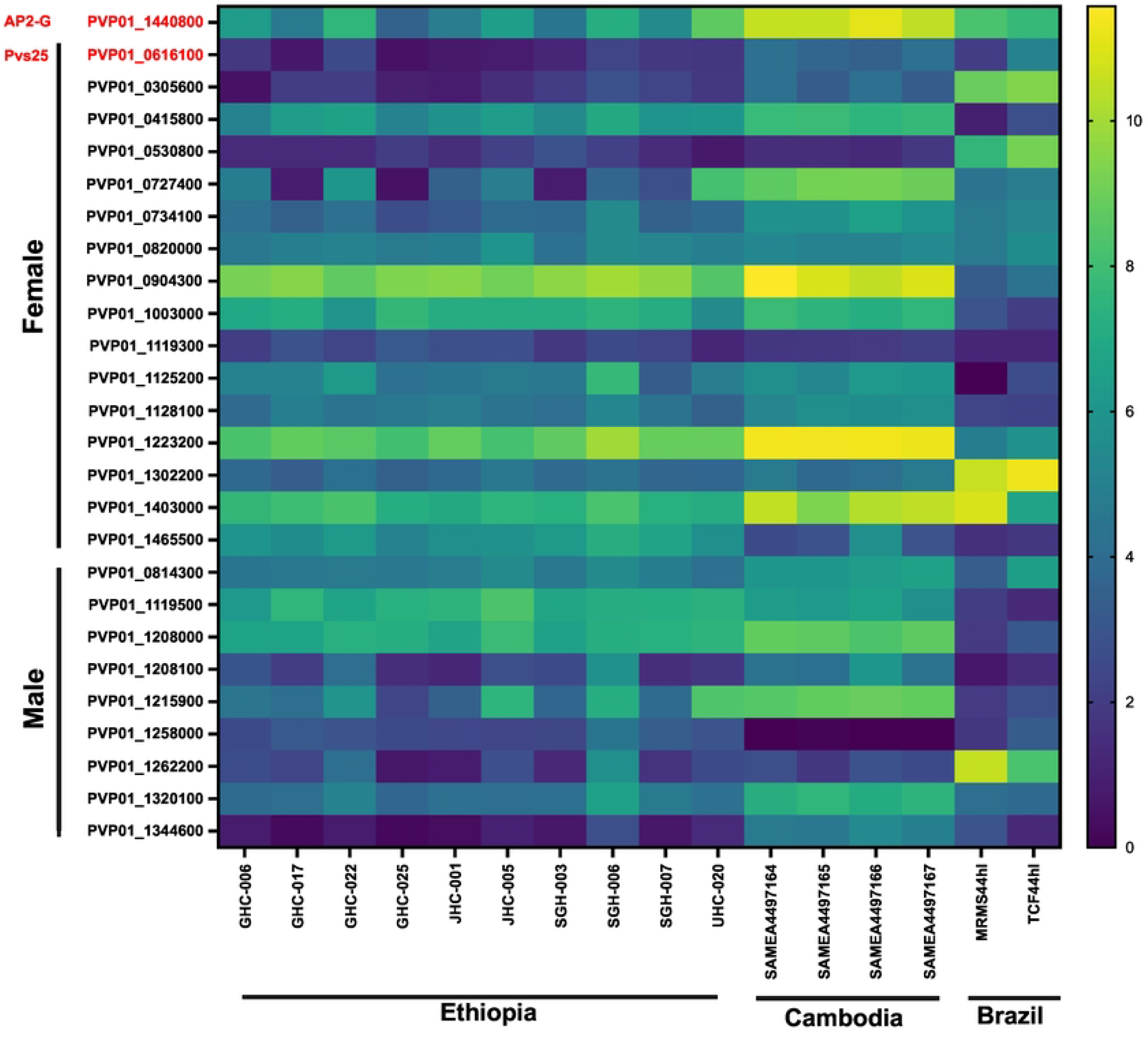
Heatmap comparing 26 *P. vivax* gametocyte biomarker candidates across the Ethiopian, Cambodian, and Brazilian *P. vivax*. Based on the expression level of *Pvs25*, all 10 *in vitro P. vivax* samples from Ethiopia, four samples from Cambodia, and two samples from Brazil contained gametocytes. Three genes, PVP01_1440800 (*PvAP2-G*), PVP01_1403000 (gametocyte associated protein, GAP), and PVP01_1208000 (*Pvs47*) from female and male gametocytes, respectively, showed the highest expression across all geographical isolates, and were consistently higher than *Pvs25*.

## 4. Discussion

This study is the first to examine the transcriptomic profile of *P. vivax* from Africa and compare gene expression among geographical isolates. Approximately 32% of the detected transcripts are of unknown function, some of which such as PVP01_0319500, PVP01_1011500, and PVP01_1228800 were among the highest expressed and could play critical function. It is not surprising that 23% of the highly expressed transcripts belong to housekeeping function, such as several zinc fingers and ATP-synthase proteins. Besides, there is a large number of highly expressed protein regulators and PTMs that have not been thoroughly examined. For example, PVP01_1444000, a ubiquitin-activating enzyme, was among the highest expressed transcripts but with unclear function. Several other protein kinases, lysophospholipases, and chaperones were also highly expressed but their role in intercellular signaling pathways is unclear. It is worth noting that a great proportion of transcripts responsible for ribosomal protein production were also highly expressed. These ribosomal proteins support intraerythrocytic development of the parasites from one stage to another.

Members of the RBP family including *PvRBP1a, PvRBP2a, PvRBP2b*, and *PvRBP3* were consistently highly expressed across the Ethiopian and Cambodian but not the Brazilian isolates, suggestive of potential differences in their role of erythrocyte invasion. Recent studies showed that the binding regions of *PvRBP1a* and *PvRBP1b* are homologous to that of *PfRh4*, and the amino acids at site ~339-599 were confirmed to interact with human reticulocytes (50).

Though the host receptors of both PvRBP1a and PvRBP1b proteins are unclear, their receptors are neuraminidase resistant (22). The PvRBP2b-TfR1 interaction plays a critical role in reticulocyte invasion in Duffy-positive infections (25). *PvRBP2d, PvRBP2e*, and *PvRBP3* are pseudogenes that share homology with other *PvRBPs* but encode for nonfunctional proteins (51). The extent to which of these *PvRBP* genes involve, if any, in erythrocyte invasion remains unclear and requires functional assays in broad samples. The high expression of *PvRBP* genes in Ethiopia could be related to a greater proportion of individuals with weak DARC expression (i.e., Duffy-negatives) (3); whereas in Cambodia and the inland regions of Brazil, populations are predominantly Duffy-positive (3). Given that *P. falciparum* can modulate gene expression in response to their hosts through epigenetic regulation (35, 52, 53), higher *PvRBP* expression in the Ethiopian *P. vivax* may allow the parasites to infect and adapt to both Duffy-positive and Duffy-negative populations (54). Further investigation on the expression and binding affinity of these *PvRBP* genes in different Duffy groups is necessary.

Another invasion protein, RIPR, was also among the highly expressed transcripts in *P. vivax*. RIPR is currently known as a vaccine target in *P. falciparum* (55), where RIPR (PfRH5) binds to the erythrocyte receptor basigin (56, 57). The PfRh5 complex is composed of PfRh5, Ripr, CyRPA, and Pf113, which collectively promote successful merozoite invasion of erythrocytes by binding to basigin (BSG, CD147) (57, 58). A BSG variant on erythrocytes, known as Ok^a-^, has been shown to reduce merozoite binding affinities and invasion efficiencies (56), though this has only been reported in individuals of Japanese ancestry (59). Despite the clear role of RIPR in *P. falciparum, P. vivax* RIPR does not seem to bind to BSG (60) and the exact role of RIPR and its binding target(s) remains unclear.

The KR-DE analysis showed 10-26% variation among the transcriptomes of the three countries, with the Ethiopian and Cambodian *P. vivax* being most similar whereas the Cambodian and Brazilian *P. vivax* most different. Genes that showed the highest levels of differentiation were those involved in housekeeping, PIR, and ribosomal functions. The exact reason for such differences amongst the geographical *P. vivax* isolates remains unclear. Previous whole genome sequencing analyses indicated that the Ethiopian, Cambodian, and Brazilian *P. vivax* are independent subpopulations, with isolates from Southeast Asia and East Africa share the most common ancestry (61). This genetic relationship may explain variations in the expression profiles. The high expression observed for some PIR proteins, such as PVP01_1000200, in the Cambodian and Ethiopian *P vivax* may suggest the prominent role of VIR antigens in epigenetic regulation associated with host exposure and immune responses (35, 52, 53), and such immune responses could vary in diverse geographical settings (62–64). Varying expression of ribosomal proteins, such as PVP01_0827400 (60S ribosomal protein L26) and PVP01_1013900 (40S ribosomal protein S9, putative) may be attributed to host nutrition, which is directly proportional to the speed of replication in *P. berghei* (65). In *P. falciparum*, host nutrition has been shown to significantly alter gene expression related to housekeeping, metabolism, replication, and invasion/transmission (65). Malnourishment has a protective effect to *P. vivax* infections in people from the western Brazilian Amazon (66). In zebra fish, sex determination can cause significant expressional differences in the housekeeping genes (67), suggesting that sexual development factors may alter expression profiles. The marked differences observed in the Brazilian isolates may also be attributed to the presence of ring stage parasites or oxidative stress related to different *in vitro* environments. Future studies should expand geographical samples of *P. vivax* and examine further host factors associated with gene expression.

In this study, the deconvolution of stage-specific transcripts was based on the *P. berghei* orthologues rather than the single-cell RNA-seq data of *P. vivax* because the latter showed little expression from the ring stage. To date, *P. berghei* remains the most comprehensively characterized single-cell data for both sexual and asexual blood stages of *Plasmodium* (68, 69), and their orthologues have been shown to be reliable for determining stage-specific transcripts (46). In primates, most *P vivax* genes have been shown to transcribe during a short period in the intraerythrocytic cycle (31) with a high proportion of late-schizont transcripts expressed as early as the trophozoite stage. In *P. berghei*, the process of gametocyte development and genes involve in sequestration are transcribed much earlier during the trophozoite-schizont transition stage. Male gametocyte development precursors are expressed in the asexual stages prior to the onset of gametocyte development (70, 71). For example, the transcription factor *AP2-G* in *P. vivax* expresses early in the asexual stage for parasites that are committed to sexual development (47). These factors hinder deconvolution efforts, making it challenging to identify precisely which genes are transcribed in each stage. Future studies should consider combining *in vivo* (rich in ring and trophozoites) and *in vitro* (rich in trophozoites and schizonts) RNA-seq data to provide a more comprehensive and reliable stage-specific model for deconvolution.

Low density *P. vivax* gametocytes in asymptomatic carriers can significantly contribute to transmission (72, 73). In areas with low transmission, submicroscopic infections are hidden reservoirs for parasites with high proportions of infectious gametocytes (74). The current gametocyte biomarkers *Pvs25* (PVP01_0616100) and *Pvs16* (PVP01_0305600) account only for female gametocytes (75), and grossly underestimate the total gametocyte densities. We previously described two alternative female (PVP01_0415800 and PVP01_0904300) and one male (PVP01_1119500) gametocyte genes that show higher expression than *Pvs25* in the Ethiopian isolates (36). Nevertheless, these genes showed relatively low expression in the Cambodian and Brazilian isolates. By contrast, *PvAP2-G* (PVP01_1440800), GAP (PVP01_1403000), and *Pvs47* (PVP01_1208000) were moderately expressed across all geographical isolates and at a level higher than *Pvs25*. These genes warrant further investigations on their potential utility as gametocyte biomarkers in low-density infections, as well as their exact role in gametocyte development.

## 5. Conclusion

This paper characterized the first *P. vivax* transcriptome from Africa and identified several host-interaction gene transcripts, including *PvRBP2a, PvMSP3.8, PvTRAg14*, and *PvTRAg22* that were highly expressed compared to *PvDBP*1 in Duffy-positive individuals. These transcripts may play prominent roles in erythrocyte invasion and merit further investigations on their binding affinity and function. We further demonstrated 10-26% differences in the gene expression profile amongst the geographical isolates, with the Ethiopian and Cambodian *P. vivax* being most similar. These findings provide an important baseline for future comparisons of *P. vivax* transcriptomes from Duffy-negative infections. Furthermore, *PvAP2-G* (PVP01_1440800), GAP (PVP01_1403000), and *Pvs47* (PVP01_1208000) of both female and male gametocytes showed higher expression than the standard *Pvs25* in all geographical *P. vivax*. These gene may provide better gametocyte detection for low-density infections.

## Acknowledgements

We thank the field team from Jimma University for their technical assistance; the communities and hospitals for their support and willingness to participate in this research; and undergraduate students at UNC Charlotte for assistance with the experiments.

## Financial support

This research was funded by National Institutes of Health R01AI162947.

## Potential conflicts of interest

The authors declare no conflict of interest.

## Supplementary Files

**Supplementary Table 1.** Name and gene ID of 43 candidate invasion genes.

**Supplementary Table 2.** Raw reads of 10 Ethiopian *P. vivax* transcriptomes in CPM and TPM metrics, Raw reads of four Cambodian *P. vivax* transcriptomes in TPM metric, and Raw reads of four Brazilian *P. vivax* transcriptomes in TPM metric.

**Supplementary Table 3.** Kenward-Roger DE analyses comparing the differentially expressed genes between (a) Ethiopian and Cambodian, (b) Ethiopian and Brazilian, and (c) Cambodian and Brazilian *P. vivax*.

## Notes

### Competing Interest Statement

The authors have declared no competing interest.

